# Risk-averse optimization of genetic circuits under uncertainty

**DOI:** 10.1101/2024.11.13.623219

**Authors:** Michal Kobiela, Diego A. Oyarzún, Michael U. Gutmann

## Abstract

Synthetic biology aims to engineer biological systems with specified functions. This requires navigating an extensive design space, which is challenging to achieve with wet lab experiments alone. To expedite the design process, mathematical modelling is typically employed to predict circuit function *in silico* ahead of implementation, which when coupled with computational optimization can be used to automatically identify promising designs. However, circuit models are inherently inaccurate which can result in sub-optimal or non-functional *in vivo* performance. To mitigate this issue, here we propose to combine Bayesian inference, Thompson sampling, and risk management to find optimal circuit designs. Our approach employs data from non-functional designs to estimate the distribution of the model parameters and then employs risk-averse optimization to select design parameters that are expected to perform well given parameter uncertainty and biomolecular noise. We illustrate the approach by designing robust adaptation circuits and genetic oscillators with a prescribed frequency. The proposed approach provides a novel methodology for the design of robust genetic circuitry.

## 1 Introduction

A key challenge in synthetic biology is the design of gene circuits with predictable functions *in vivo*. Common approaches to circuit design often rely on mathematical models coupled with optimization algorithms to find suitable designs *in silico*. Recent examples include the use of gradient-based or Bayesian optimization [11, 19], as well as exhaustive search [14, 16, 24], mixed-integer optimization [6, 22], and multiobjective optimization [32]. While these approaches can accelerate the design process, their effectiveness is limited by the inaccuracies of the *in silico* models, as these limit their ability to predict circuit function *in vivo*. Consequently, a circuit design may work as intended *in silico* but not in vivo.

Model inaccuracies can result from molecular interactions that are unknown or unaccounted for, or from uncertainty in model parameters such as kinetic binding rates or dissociation constants. These parameters are challenging to estimate accurately as they typically require bespoke experimental work solely for the purpose of model estimation. To address this problem, early work by Barnes et al [4] and Woods et al [33] employed Approximate Bayesian Computation to quantify the biological robustness of various circuits under uncertainty, while Schladt et al [28] considered the design of transcriptional logic gates. Recent work by Sequeiros et al [29] modelled noisy gene expression with Partial Integro-Differential Equations and optimized circuit function with mixed-integer programming. In this work, we introduce tools from uncertainty quantification and risk management for the optimization of genetic circuits. Our approach allows us to leverage experimental *in vivo* data from failed prototypes to propose new designs that are more likely to function as intended.

In the fields of uncertainty quantification and decision theory, model uncertainty is broken into two components: *epistemic* and *aleatoric* uncertainty [13]. Epistemic uncertainty is caused by limited system knowledge and translates into uncertainty on model parameters. Aleatoric uncertainty, on the other hand, is caused by the intrinsically stochastic nature of biochemical processes [8]. This distinction is important because epistemic uncertainty can be reduced by increasing the size of observed data, while aleatoric uncertainty is irreducible and independent of the number of experimental observations. Models with misspecified parameters can incorrectly predict system behaviour and thus guide the optimization process to sub-optimal designs (1a–b), while biochemical stochasticity can change the intended function of gene circuits [8].

Here, we present a strategy for gene circuit optimization that jointly accounts for biochemical stochasticity and parameter uncertainty, using a combination of Bayesian inference and tools from risk management [5, 20, 21]. Our approach relies on the distinction between design parameters, i.e. those that designers can control with genetic parts such as promoters, termination sites, or degradation tags, from uncertain parameters, i.e. those parameters that are challenging to modify experimentally and are not known *a priori* with sufficient certainty. Employing data from preliminary, possibly non-functional designs, we first quantify parameter uncertainty with Bayesian inference and then employ risk-averse optimization to select design parameters that are expected to perform well under the estimated posterior distribution and biomolecular noise.

More technically, after estimating the posterior distribution of the uncertain parameters given the available data, we employ *Thompson sampling* [31] to obtain a set of promising designs. Thompson sampling is a commonly used decision-making procedure that draws samples from the distribution of possible optimal designs. This can be done by drawing a sample from the posterior distribution of the uncertain parameters and then minimizing the expected loss with respect to the design parameters [9]. By repeating this procedure with different posterior samples we find sets of promising designs. For each design, we determine the predictive performance distribution reflecting both the epistemic uncertainty of the parameters (via their posterior distribution) and biomolecular stochasticity. We summarise the predictive performance distributions in terms of expected performance and risk, and, to avoid critical circuit failures, we select risk-averse designs, i.e. designs which produce desired outcomes even under unfavourable conditions (Figure 1c–d).

**Fig. 1.**
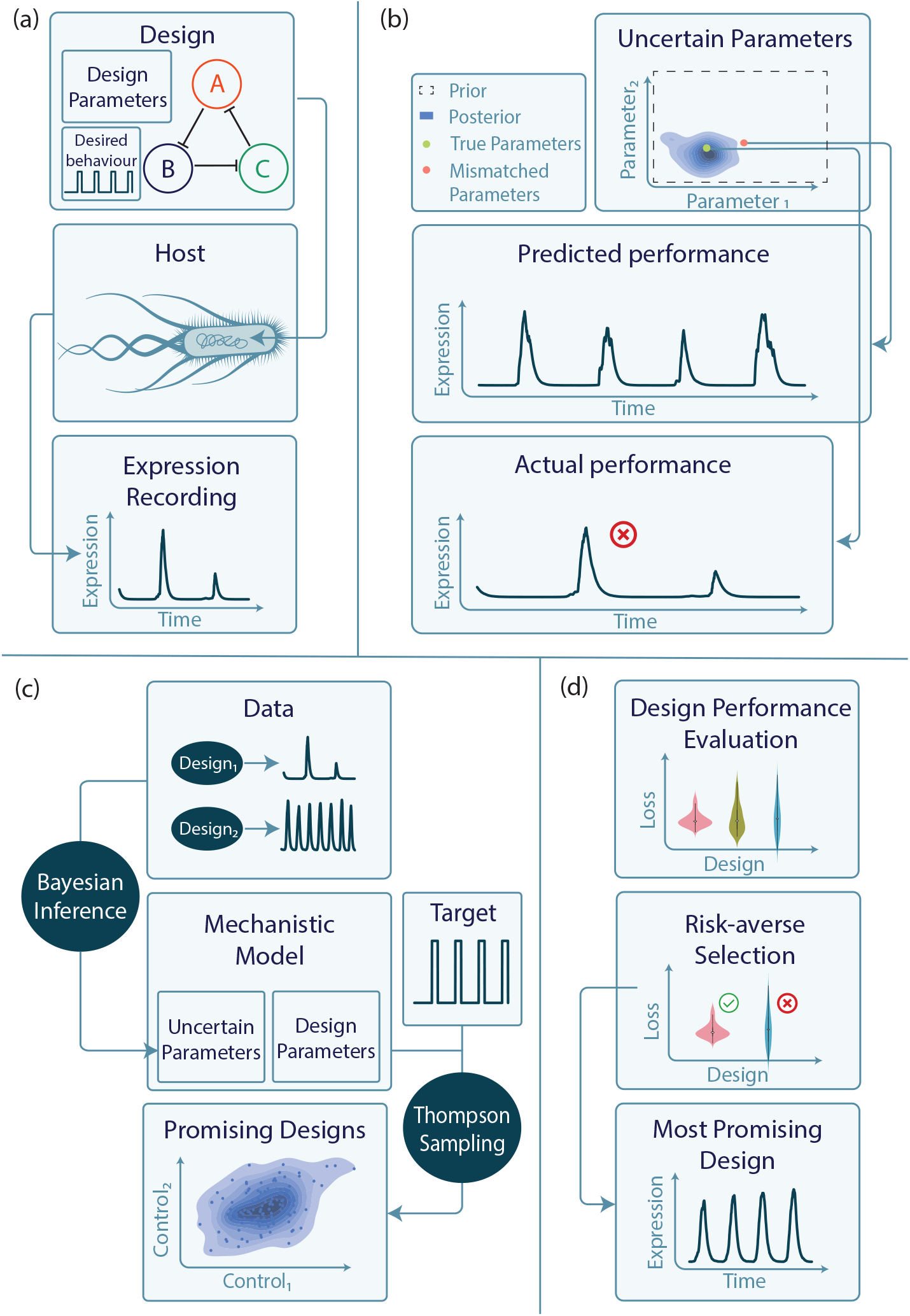
**(a)** Introducing specific genetic design into the host organism results in the system performing specific functions. During the experiment, the expression levels can be recorded and used as data for inference. The designs are controlled by the design parameters, with the objective of creating an oscillator that produces four pulses within a designated time frame. **(b)** The lack of knowledge about the uncertain parameters can lead to wrong design choices. In particular, a design can achieve reasonable performance under mismatched parameters (here: 4 oscillations), but not when deployed under the true parameters. **(c)** To solve the problem introduced in (b) we propose to use the data observed in previous experiments to infer the uncertain parameters of the mechanistic model via Bayesian inference. The uncertainty about the parameters causes uncertainty on the level of the optimal designs, which we capture with *Thompson sampling*. **(d)** We evaluate the performance of the acquired designs and use risk-averse selection to pick the design(s) that are most likely to achieve the desired function even under unfavourable conditions.

We illustrate the approach with two exemplar design tasks: circuits that achieve robust perfect adaption and circuits that produce oscillatory dynamics with a prescribed frequency. To the best of our knowledge, this is the first application of risk-averse optimization for gene circuit design under uncertainty and lays the groundwork for improved computational strategies for designing circuits with predictable functionality.

## 2 Results

We test our method on the design of a perfect adaptation circuit [30] and genetic oscillator (repressilator) [7]. We use synthetic data to evaluate the method against known ground truth, using large observational noise or stochastic models to closely emulate biological measurements encountered in practice. We demonstrate that the proposed framework is compatible with different approaches to Bayesian inference and Thompson Sampling: for the perfect adaptation circuit, we use Markov Chain Monte Carlo (MCMC) [12] for inference and stochastic gradient descent to acquire the Thompson samples. For the design of the genetic oscillator, we use Sequential Monte Carlo Approximate Bayesian Computation (SMC-ABC) for inference, see e.g. [15], and Bayesian optimization [9] to obtain the Thompson samples.

### 2.1 Perfect Adaptation

In this case study, we design a simple robust perfect adaptation (RPA) circuit [30]. RPA systems return to their previous steady state after persistent changes in their inputs. Such functionality is important as it ensures that genetic circuits can sustain their function in the face of external perturbations.

### 2.1.1 Inference

We used a circuit which comprises two genes, as shown in Figure 2a. Their dynamics are modelled by the following Ordinary Differential Equations (ODEs) [30]:

**Fig. 2.**
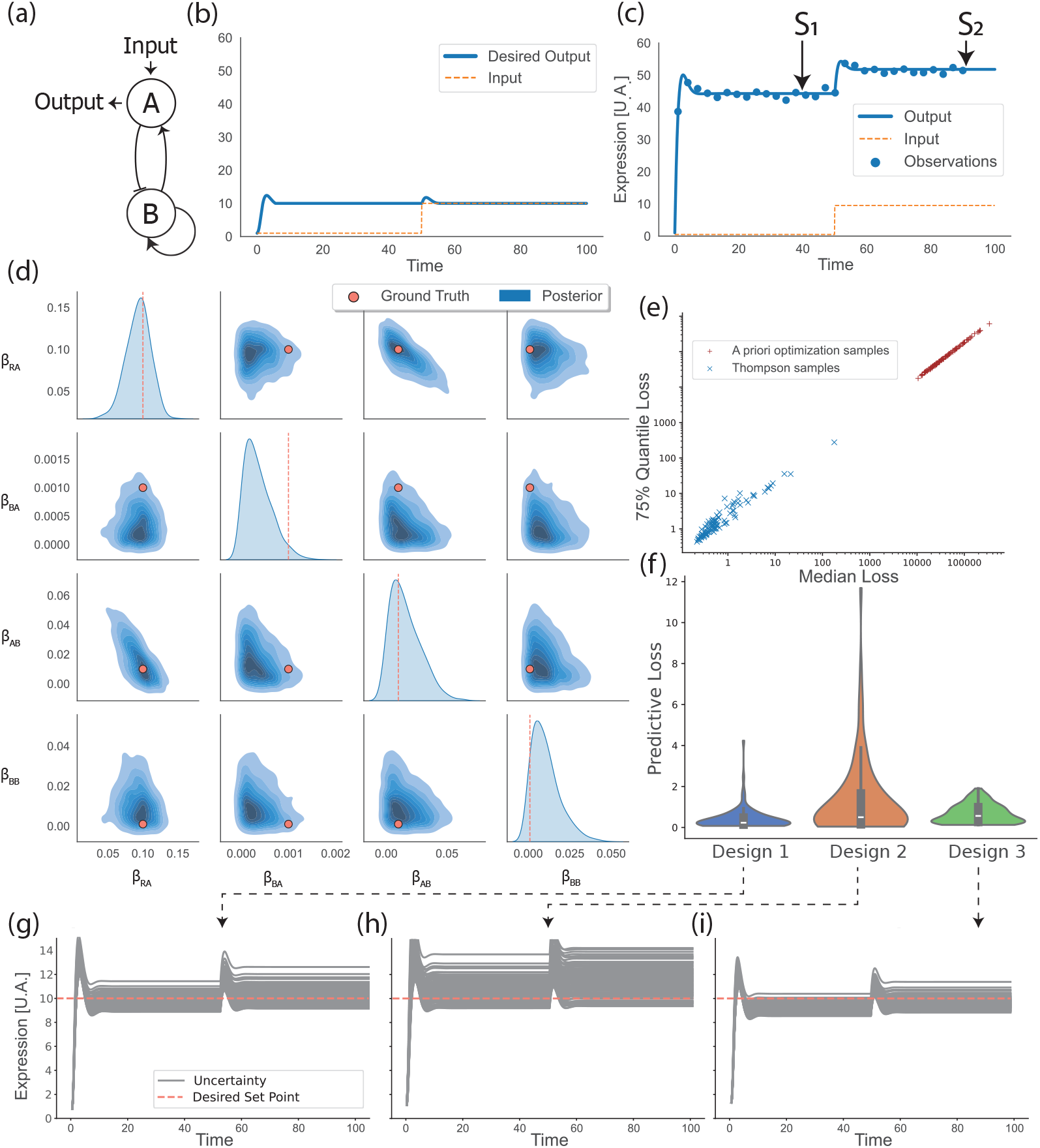
Design of a robust perfect adaptation (RPA) circuit. (a) Schematic representation of the circuit. (b) The desired behaviour of the circuit adapting to an input change with a set point of 10 U.A. (c) Observed data from a failed design that does not adapt to the change of the input. (d) Posterior over uncertain parameters. (e) Performance of Thompson samples contrasted with *a priori* optimization samples. (f) Performance of three representative designs (Thompson samples). (g) Predictive performance of Design 1 (h) Predictive performance of Design 2 (note larger uncertainty corresponds to larger spread of predicted trajectories compared to Design 1). (i) Predictive performance of the Design 3 (lowest spread but achieved equilibrium tends to be worse than Design 1).

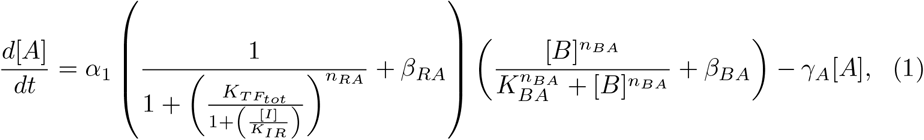

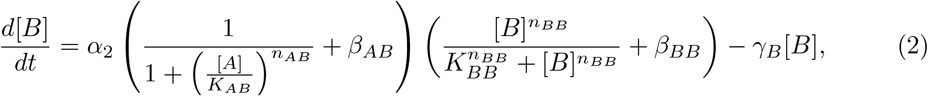

where *I* is an external input, *α*_*i*_ is the maximal expression level and *γ*_*i*_ is the degradation rate for the proteins encoded by genes *i* ∈ {*A, B*}. The parameters *K*_*ij*_, *n*_*ij*_ and *β*_*ij*_ are the regulatory thresholds, Hill coefficients, and activation/repression leakages from node *i* to *j*, while *K*_*IR*_ and *K*_*T*_ and 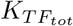 are affinity constants of input activation to node A. Further, *n*_*RA*_ and *β*_*RA*_ are a Hill coefficient and the leakage of the activation from the input to node A. For a fixed concentration of the input *I*, the system has a single stable equilibrium point, and our design goal is to find circuit parameters for which such equilibrium is equal to 10 U.A. insensitive to step changes in the input, as shown in Figure 2b.

We assume that the controllable design parameters are the maximal expression levels and regulatory thresholds, *ϕ* = (*α*_1_, *α*_2_, *K*_*BA*_, *K*_*AB*_, *K*_*BB*_), as these are typically amenable for experimental tuning with DNA sequence engineering [2, 18]. The uncertain parameters are the leaky expression levels of each protein, *η* = (*β*_*RA*_, *β*_*BA*_, *β*_*AB*_, *β*_*BB*_), for which we assume uniform priors *U* (0, 1). We chose leakages as the uncertain variables as they are more challenging to manipulate experimentally [3] and negatively affect system performance [23]. The rest of the parameters were fixed to values provided in the Supplementary Material.

To obtain observed data, we simulated the ODEs with a ground truth value of the uncertain parameters equal to *η*_*GT*_ = (0.1, 0.001, 0.01, 0.001) and the design parameters set to *ϕ*_1_ = (1.0, 1.0, 1.0, 1.0, 1.0); the design parameters *ϕ*_1_ were deliberately chosen to represent a failed design attempt where the system reaches a new equilibrium after a step change in the input signal. To emulate real-world data, we added observational noise and assumed a relatively small number of temporal measurements (*N* = 30 as shown in Figure 2c).

We used the observed data from the failed design to infer the uncertain parameters using the Markov Chain Monte Carlo NUTS sampler [12]. We ran three independent chains of 1000 samples (with 500 burn-in samples), all resulting in similar results indicating successful mixing (Supplementary Figure S4). The posterior distribution is shown in Figure 2d. The ground truth values *η*_*GT*_ are contained in the main bulk of the posterior probability mass but they are at times not equal to the posterior mode or mean (e.g. *β*_*BA*_), which is normal in Bayesian inference when the amount of data is not large. Importantly, the maximum likelihood estimate (i.e. the mode of the posterior as the prior is uniform) does not coincide with ground truth, and, if used in design, would lead to circuits that may not work well when deployed. In contrast, as we show next, our approach considers the whole distribution of uncertain parameters; as a result, the correct ground truth model will be considered for design and, moreover, performing risk-averse design becomes possible, resulting in circuits that work well even in unfavourable conditions.

#### 2.1.2 Design

To find promising designs, we acquired 1000 Thompson samples of the design parameters. The optimization loss for a Thompson sample was

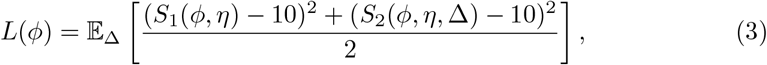

where *η* is a fixed posterior sample, *S*_1_(*ϕ, η*) is the steady state achieved when the input equals *I* = 1 before any change and *S*_2_(*ϕ, η*, Δ) is the steady state achieved after the input was changed to *I* = 1 + Δ where Δ was drawn from a uniform distribution on (1, 10). Minimising the loss function with respect to the design parameter *ϕ* ensures that the steady state both before and after the input change is close to the desired value of 10 U.A. for a fixed posterior sample *η*. We randomised the input change to ensure that the steady state output is close to the target across the whole range of potential input concentration changes. Note that, the observational noise does not feature in the loss function as it does not impact the Thompson sampling (see S3 in Supplementary Material). The optimization with respect to the design parameters *ϕ* was performed using stochastic gradient descent with automatic differentiation. By repeating the minimisation for 1000 posterior samples *ϕ* we acquired a batch of 1000 promising designs. Figure 3b shows their distribution.

**Fig. 3.**
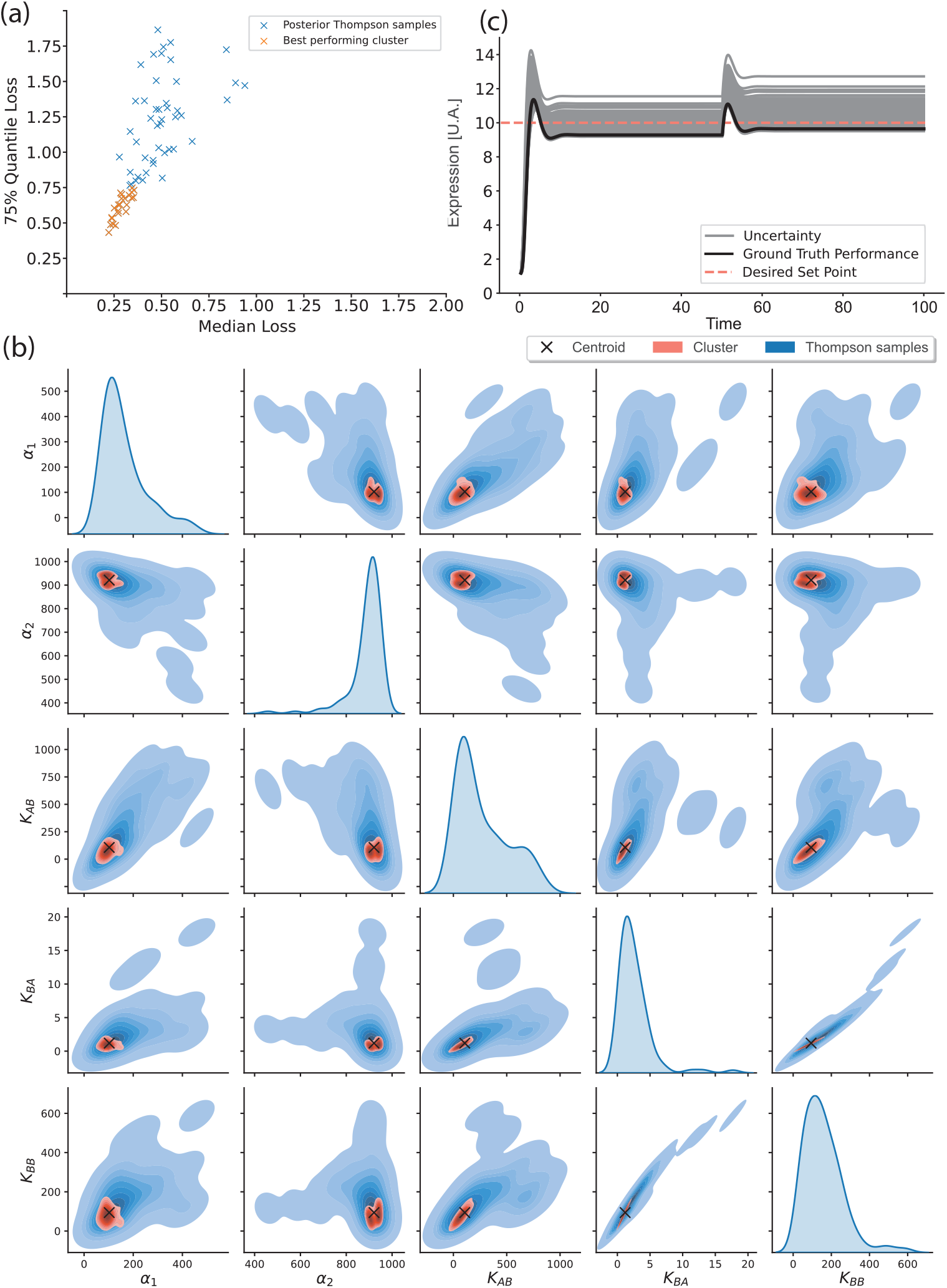
Design of a robust perfect adaptation (RPA) circuit. (a) Median and 75%-quantile of the loss distribution of each Thompson sample with highlighted best-performing designs. The cut-offs are 0.5 and 0.75 for median and 75%-quantile loss respectively. (b) All the Thompson samples together with the best-performing ones in the design space. The best performing designs turn out to form a cluster. (c) The predictive behaviour of the centroid of the best-performing cluster. A solid black curve shows the performance under ground truth parameters *η*_*GT*_, while grey trajectories present the performance for multiple posterior samples, thus displaying the epistemic uncertainty.

#### 2.1.3 Evaluation

To evaluate the quality of the designs we computed the predictive loss distribution *p*(*L*|*ϕ, 𝒟*_*o*_) for each acquired design *ϕ* (see Methods), where 𝒟_*o*_ is the observed data. The predictive loss distribution quantifies the performance of a design reflecting both the epistemic uncertainty about the model parameters and biomolecular stochasticity. To assess the tradeoffs of deploying different designs, Figure 2f shows the predictive loss distributions for three exemplar designs. The first design has a lower median loss (middle dot of the violin) than the second one but the worst-case loss is higher. On the other hand, the third design has both higher median and worst-case loss than the other two, but lower best-case loss. Depending on the metric and the designer’s preference, different designs might be preferred: Design 1 is the best according to the median loss, Design 2 according to the best-case loss and Design 3 according to the worst-case loss.

The shape of the predicted loss distribution is further reflected in Figure 2g–h where the predictive behaviour of the first, second and third designs is shown for multiple posterior samples of uncertain parameters (multiple grey trajectories). The predicted behaviour of the first design is more concentrated around the desired set point compared to the second design, which reflects the shorter upper tail of the first design’s violin plot. The third design usually results in a lower equilibrium than desired which is reflected in the worse median performance compared to the first design.

For the purposes of robust design, we reasoned that using the worst-case loss as risk-averse metric is not the best choice if the worst-case outcome is rather unlikely. Instead, we opted to use a less pessimistic but still risk-averse 75%-quantile to rank the designs, together with the median of the predictive loss distribution. The median robustly measures the expected predicted outcome, while the 75%-quantile indicates how much worse the loss is predicted to get if performing worse than the median (see Methods). We thus rank the designs based on their predicted performance (measured by the median) and their safety (assessed by the 75% quantile). Figure 2e plots the two metrics against each other for the designs learned by Thompson sampling (blue crosses). We see that we do not need to compromise between performance and safety; the two metrics are here positively correlated with each other.

Our method uses Bayesian inference to reduce the prior uncertainty, promising better designs acquired under posterior (reduced) uncertainty. To verify the efficacy of the method, Figure 2e further compares the Thompson samples (blue crosses) with the prior optimization samples (red “+” signs) obtained by the same procedure, but using the prior rather than the posterior distribution for the uncertain parameters. The Thompson samples improve over the prior optimization samples by an order of magnitude, both in terms of median and 75% quantile loss. This demonstrates that by reducing uncertainty with observed data, we can acquire better designs, which is the premise of our work.

While we acquired a batch of promising designs, it may only be possible to test a single or few designs experimentally. To locate the best-performing design, we first filter the designs with the best median and 75%-quantile performance in Figure 2e, choosing cut-off values of 0.5 for the median and 0.75 for the 75%-quantile loss. Figure 3a highlights the selected designs. These best-performing designs turn out to form a cluster in the design space as shown in Figure 3b. We used the centroid of this cluster to pick the single most promising design. Figure 3b shows that this centroid design achieves a predictive performance (grey trajectories) close to the desired 10 U.A., as well as good performance when deployed in the system with the ground truth parameter values *η*_*GT*_ (solid black trajectory). This demonstrates that the proposed approach was effective in designing a RPA circuit under uncertainty.

### 2.2 Repressilator

To test our methodology in circuits with more complex dynamics, we focussed on the repressilator, a landmark system in synthetic biology [7] owing to its ability to produce temporal oscillations in protein expression levels.

#### 2.2.1 Inference

In this case study, we use a stochastic model of the Repressilator [7] parameterized by *θ* = (*α, n, K, γ*), which represents maximal expression, Hill coefficient,regulatory threshold, and degradation rate, respectively. The stochastic differential equations for each protein *p*_*i*_, *i* ∈ {1, 2, 3}, are

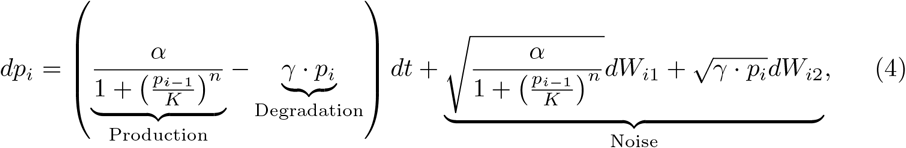

where *p*_0_ := *p*_3_ and *W*_*ij*_ is an independent Wiener Processes for *i* ∈ {1, 2, 3}, *j* ∈ {1, 2}. A schematic representation of the circuit is shown in Figure 4a.

**Fig. 4.**
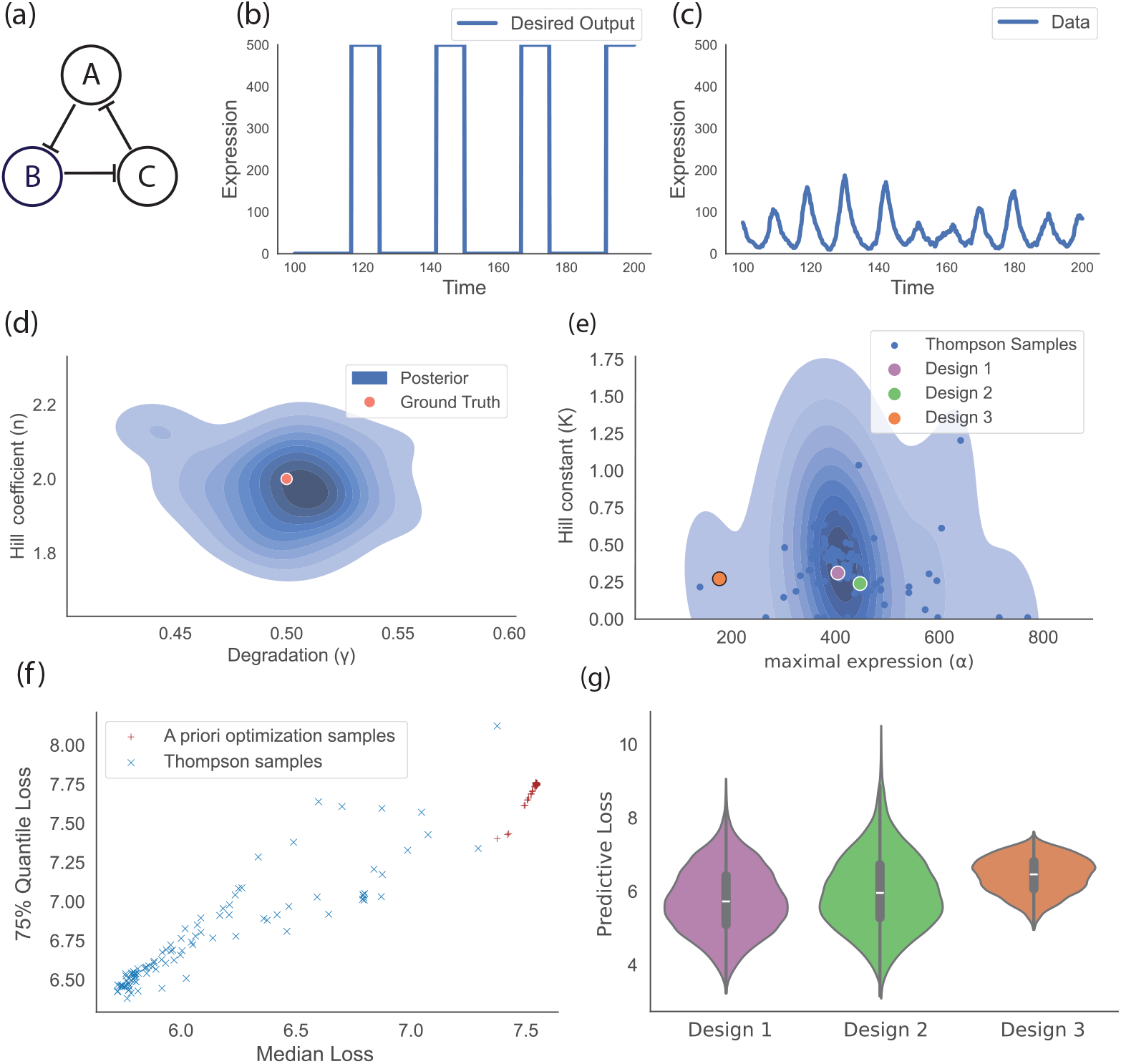
Design of a repressilator. (a) Schematic representation of the circuit. The design parameters are maximal expression rate *α*, and regulatory threshold *K*. The uncertain parameters are Hill coefficient n and degradation rate *γ*. (b) Observed data from a failed design that does not result in 4 oscillations in the 100-200 min timeframe. (c) The desired behaviour of the circuit. (d) Posterior over uncertain parameters. (e) The designs acquired with Thompson sampling together with three representative ones used for further evaluation. (f) The evaluation of the Thompson samples (blue crosses) contrasted with *a priori* optimization samples (red “+” signs). Note that quantifying the posterior significantly improves the performance of the acquired designs. (g) Predictive performance of three representative designs.

The Hill coefficient *h* and degradation rate *γ* are the uncertain parameters *η* = (*h, γ*), for which we assumed a uniform prior distribution *U* (0, 10). The maximal expression *α* and regulatory threshold *K* are the design parameters *ϕ* = (*α, K*). The goal in this case study is to find a design that produces 4 oscillations in a timeframe (100, 200) min, Figure 4b displays the desired behaviour of the system.

To create the observed data we simulated a trajectory with design parameters *ϕ*_1_ = (334, 10) and ground-truth values of the uncertain parameters equal to *η*_*GT*_ = (0.5, 2.0). Figure 4c shows the obtained trajectory. The design *ϕ*_1_ fails to produce the desired result, portraying an unsuccessful design attempt as it exhibits 10 peaks rather than 4 in the specified timeframe. Rather than discarding the failed attempt, our approach uses it to learn the uncertain parameters and quantify their uncertainty, which we then further use for risk-averse optimization.

We inferred the uncertain parameters from the observed data using SMC-ABC [e.g., 15], which is a Bayesian inference approach for models with intractable likelihoods, which is here the case due to the SDE model in (4). The method matches the observed data with data simulated from the model, using a distance metric to measure the difference. Due to the stochastic nature of the system, each time the SDEs are solved, the oscillations have a different phase. Therefore we developed a phase-independent distance metric that operates on the spectrum of the Discrete Fourier Transform (DFT). Denoting by *F*_(*η,ϕ*)_ and *F*_*obs*_ the discrete Fourier transforms of the solution *f*_(*η,ϕ*)_ of the SDE and the observed signal *f*_*obs*_, respectively, the distance metric is:

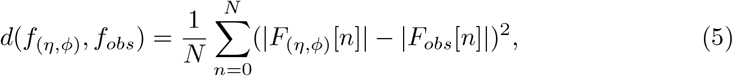

where *N* is the length of the discrete Fourier transform. Figure 4d shows the posterior distribution of the uncertain parameters inferred with SMC-ABC. Note that the ground truth parameters *η*_*GT*_ are contained within the posterior distribution.

#### 2.2.2 Design

To find the promising designs under the uncertainty, we acquired 100 Thompson samples using Bayesian Optimization. The objective was to match the model’s output with the desired target shown in Figure 4c. As in the inference case, we used a phase-independent approach to match the SDEs solution *f*_(*η,ϕ*)_ to the target *f*_*target*_. Denoting by *F*_(*η,ϕ*)_ and *F*_*target*_ the DFT of those signals, the objective was to minimize

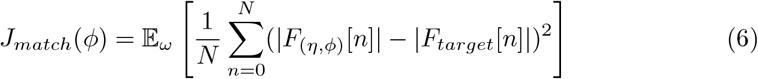

with respect to *ϕ* while *η* was kept fixed to a posterior sample. The expectation is taken over the stochasticity *ω* of the SDE model, i.e. the biomolecular noise. This was performed for 100 posterior samples, resulting in 100 Thompson samples, which are visualised in Figure 4e.

#### 2.2.3 Evaluation

We assessed the performance of the acquired designs with the predictive loss distribution *p*(*L*|*ϕ*, 𝒟_*o*_) for each design *ϕ* (see Methods). Figure 4g displays the predictive loss distribution of three exemplar designs with violin plots. We picked two designs close to the mode of the design distribution (Designs 1 and 2 in Figure 4e) and one close to the boundary (Design 3) to contrast them with each other. Design 1 and Design 2 have both better median and 75%-quantile performance than Design 3. However, comparing the lower tails of the distributions, the worst-case performance of Design 3 is better than the other two. Hence, depending on the risk-averseness, Design 1 or Design 3 may be preferable. In particular, if only the worst-case scenario is taken into account, Design 3 may be more suitable than Design 1. Here, as in the previous case study, we used the median and 75% quantile of predictive loss distribution for the selection of the designs.

We verified the efficacy of the method by comparing the acquired designs (Thompson samples), with prior optimization samples obtained by the same procedure, but using the prior rather than the posterior distribution for the uncertain parameters. Figure 4f shows the median and 75% quantile of the predictive loss distribution for both the acquired designs (Thompson samples) and the *a priori* optimization samples. The Thompson samples improve over the prior optimization samples both in terms of the median and 75% quantile loss. This shows that making use of even unsuccessful design attempts to infer the uncertain parameters, i.e. reducing their epistemic uncertainty, helps to find better designs.

Similarly to the previous case study, it turns out we do not need to compromise between expected performance and risk aversion as seen in Figure 4f. We thus pick the designs that achieve the best performance for both metrics, highlighting them in Figure 5a. The selected designs are further visualised in the design parameter space in Figure 5b. They form a clear cluster with the centroid indicated by the green dot. The centroid corresponds to the most promising design if only a single design can be tested experimentally.

**Fig. 5.**
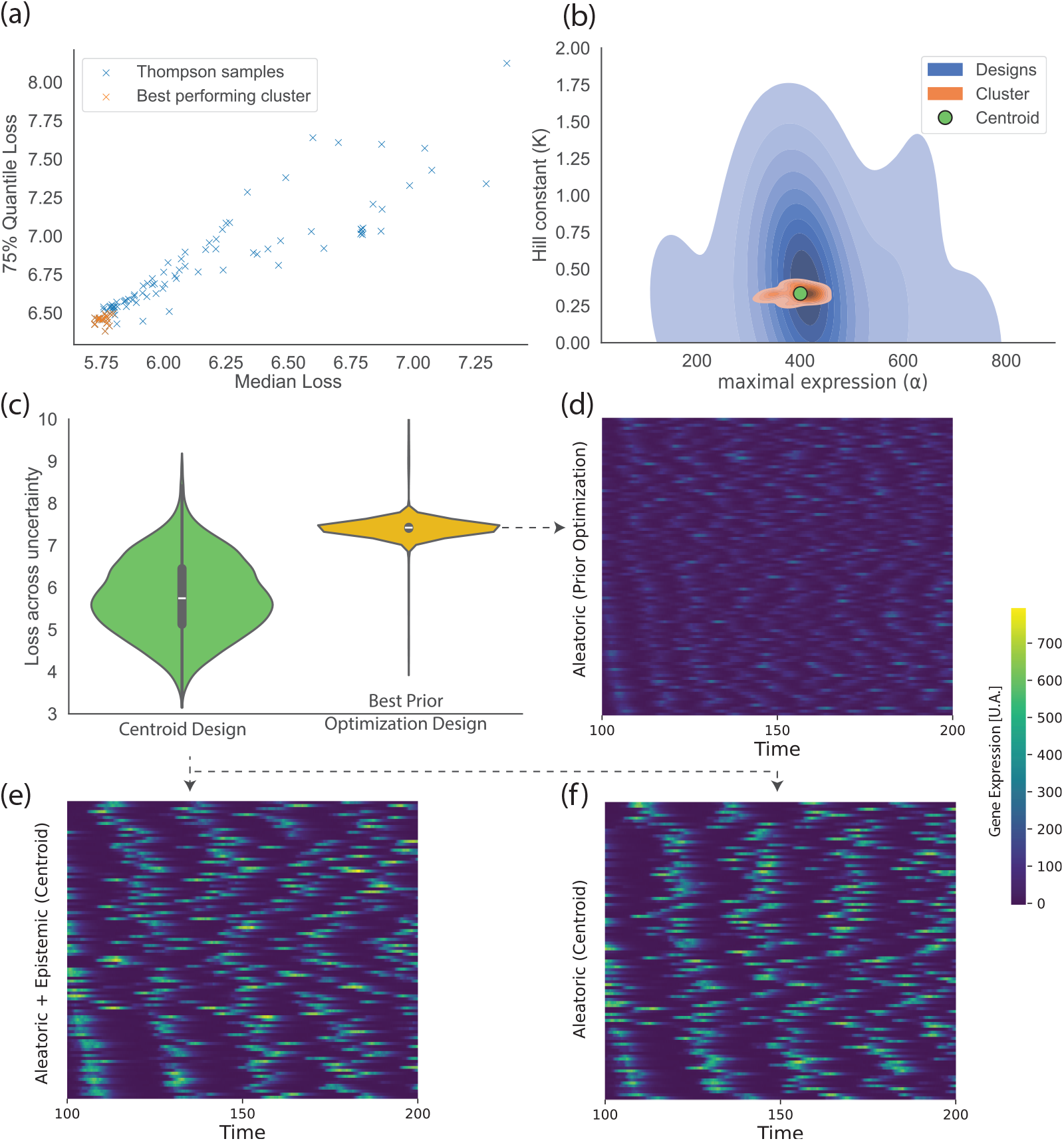
Design of a repressilator. (a) Median and 75%-quantile of the loss distribution of each Thompson sample with highlighted best-performing designs. (b) Best-performing designs form a cluster in the design parameter space. (c) The performance of the centroid of the best-performing designs across uncertain parameters together with prior optimization design with the lowest expected loss. (d) The performance of the best prior optimization design under ground truth parameters. Each row is the simulated gene expression for this design given ground truth *η*_GT_ Note that the desired 4 oscillations are not generally achieved. (e) Posterior predictive of the centroid design, the 4 oscillations are generally achieved. Each row is the simulated gene expression for this design for different posterior samples of uncertain parameters. (f) Performance of the centroid design under ground truth uncertain parameters. The design goal of four oscillations in the specified time-interval is achieved.

To evaluate the performance of the most promising design, we show in Figure 5c the predictive loss distribution of the centroid design. For reference, we also show the loss of the best-performing prior optimization sample (according to expected loss). The predictive loss distribution is shifted to smaller values for the centroid design, reflecting the benefit of reducing the epistemic uncertainty of the parameters, in line with Figure 4f.

Figure 5e visualises the posterior predictive behaviour of the repressilator for the centroid design. Each row of the heatmap is a simulated gene expression by the SDE system, using a different sample from the posterior distribution of the uncertain parameters and a different random seed *ω*, reflecting both epistemic and aleatoric uncertainty. The centroid design is predicted to yield four oscillations across different levels of uncertainty.

To emulate the scenario where the designed circuits are deployed in vivo, Figure 5d–f visualises the performance of the centroid design and the best performing prior optimization sample under ground truth parameters *η*_*GT*_. In contrast to Figure 5e, the randomness here is only due to the inherent randomness of the SDEs (aleatoric uncertainty), modelling biomolecular noise. The centroid design results in four oscillations as desired, while the best prior optimization sample fails to achieve the objective.

This demonstrates that reducing the uncertainty about the model parameters with the data from the failed design leads to better designs.

## 3 Discussion

Designing circuits *in silico* so that they have predictable functionality *in vivo* is a major challenge in synthetic biology. In this paper, we introduced a new method that combines uncertainty quantification and risk management for the risk-averse optimization of genetic circuits. The method identifies designs that are likely to perform well even under unfavourable conditions, thereby increasing the predictability of *in silico* circuit design.

The results demonstrate that our method can utilise data from failed design attempts to suggest more promising designs. Reducing the epistemic uncertainty using observed data resulted in Thompson samples with better predictive performance compared to prior optimization samples, as measured with risk-neutral and risk-averse metrics. Further, we only used a single failed experiment data to acquire well-performing designs, indicating data efficiency.

While our method has not yet been evaluated on real-life data from wet lab experiments, we present promising results using realistic synthetic data—we used a stochastic model, which mimics the noise observed in biological systems and added observational noise. A benefit of using synthetic data is that we know the ground truth parameters, which allowed us to verify the quality of the Bayesian inference process and the obtained designs.

A noteworthy future direction is addressing model mismatch. While this work addresses the issue of mismatched or uncertain parameters, the model itself can also be too simplistic. It may be that no combination of parameter values can sufficiently well represent reality. For example, this could happen when the metabolism of the organisms is omitted or the reactions between different parameters of the model are not accounted for, e.g. the repression leakage could change when a different promoter is used. Optimization of genetic circuits under this kind of model mismatch is important future work.

Our approach can be loosely seen as single-step biology-informed Bayesian Optimization (BO) [9]. While standard BO employs general-purpose models such as Gaussian Processes or ensembles of random forests, we use systems of differential equations that are mechanistic models of the biomolecular reactions. Albornoz et al [1] utilised standard BO and Radivojević et al [26] used a data-driven approach related to Bayesian optimization in the context of synthetic biology. While such purely datadriven approaches are preferred if no suitable mechanistic model is available, they require more observed data than when prior knowledge in the form of a mechanistic model is available. Indeed, our method was effective in our case studies with data from only a single failed design, demonstrating the effectiveness of using prior information in the form of mechanistic models.

Further contrasting our approach with standard BO, we note that our approach uses expression trajectories (vectors in R_*n*_) as the observed data instead of objective function values (scalars) as typically the case in standard BO. Avoiding to reduce vectors to scalars allows us to make use of all available data.

While we specifically focus on synthetic biology, our work relates to broader areas of research regarding optimization under uncertainty and risk management. For general dynamical systems, Gerlach et al [10] utilised Koopman operators to efficiently compute the expectation of the loss over the posterior which can be used as an optimization objective. In our approach, we use Thompson sampling rather than calculating the expectation, which allows us to acquire a batch of designs, rather than a single design. Batching can be useful if multiple experiments can be performed concurrently. Moreover, it leads to a distribution over optimal designs which allows us to use risk-averse measures to pick the most promising design(s), in contrast to the expectation which exhibits risk-neutral behaviour [9]. Risk-averse optimization was explored mostly in portfolio theory. It normally involves using risk-averse measures that asymmetrically summarize the performance distribution [9] such as superquantiles (conditional value-at-risk), expectiles and quantiles (value-at-risk) [5, 20, 21], which we use in our work. Finally, we note that due to the use of risk-averse measures, our approach can be seen as an instance of *robust optimization*, in the sense of Garnett [9, Chapter 11].

In conclusion, we developed an uncertainty-aware model-based design framework that makes maximal use of available data, including data from potentially unsuccessful designs. Our approach leverages risk management to obtain designs that perform well even under unfavourable conditions caused by different sources of uncertainty, both epistemic and aleatoric. As we used Thompson’s sampling to acquire designs, our approach likely can be used sequentially, balancing exploration and exploitation [9]. Such a sequential approach could allow for the continuous design loop of successive design acquisition and model updates. We envision our approach to form a basis for further research bridging the gap between *in silico* predictions and *in vivo* behaviour, aiming to acquire designs performing well not only when predicted *in silico* but also when deployed in vivo.

## 4 Methods

### 4.1 Inference

We assume that a mechanistic model of the genetic circuit is available and parameterized by *θ*. The parameters *θ* = (*η, ϕ*), are split into two subsets—the uncertain parameters *η* and the design parameters *ϕ*. Set *ϕ* are the controllable parameters that we wish to optimize to achieve the desired functionality. Parameters *η* are unknown or uncertain, but which can be inferred from observed data. The observed data 𝒟_*o*_ comprises observed trajectories *t*_*i*_ of gene expression over time and their corresponding design parameters *ϕ*_*i*_, where *i* ∈ 1, 2, …, *n* and *n* is the size of the data set. Our goal is to use the observed data together with the mechanistic model to find the most promising design parameters to achieve a desired target functionality.

We place a prior distribution *p*(*η*) over the uncertain parameters. In this work, we use a uniform prior encapsulating a reasonable range of uncertain parameters, however, a more informative prior distribution could be used, reflecting expert knowledge or previous work. Then, we use Bayesian parameter inference to update the prior belief about *η* given 𝒟_*o*_, obtaining the posterior distribution *p*(*η*|𝒟_*o*_) over the uncertain parameters. Depending on the model, we use MCMC [12] or approximate Bayesian computation [ABC, e.g. 15] to sample from the posterior distribution. We denote the set of posterior samples by 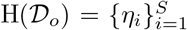 where *S* is the number of samples and *η*_*i*_ ∼ *p*(*η*|𝒟_*o*_).

### 4.2 Design

For each sample *η*_*i*_ in H(𝒟_*o*_), we optimize the design parameters *ϕ* to form the set of Thompson samples 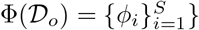, where each *ϕ*_*i*_ is the solution to an optimization problem that involves *η*_*i*_,

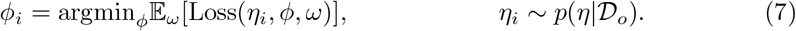

Here, *ω* represents the inherent randomness in the model, leading to aleatoric uncertainty, and “Loss” is a suitable loss function for desired functionality. Since the model produces a time-series given (*η, ϕ*) for a specific random seed *ω*, the loss, which assesses whether the generated time-series satisfies the desired functionality, depends on (*η, ϕ, ω*). We average the loss by taking the expectation over the aleatoric uncertainty *ω*, which is common practice in Thompson sampling [27].

### 4.3 Evaluation

For a fixed value of the design parameter *ϕ*, Loss(*η, ϕ, ω*) is a random variable *L* whose distribution is induced by the distribution of *η* and by *ω*. For *η* ∼ *p*(*η*|𝒟_*o*_), we denote the distribution associated with the random variable *L* by *p*(*L*|*ϕ, 𝒟*_*o*_). It represents the predictive performance of design *ϕ* under both epistemic uncertainty, via *η* ∼ *p*(*η*|𝒟_*o*_), and aleatoric uncertainty (biomolecular noise), via *ω* ∼ *p*(*ω*).

We summarise the predictive performance distribution *p*(*L*|*ϕ, 𝒟*_*o*_) via quantiles: the 50% quantile (median) as a robust, risk-neutral, measure of expected performance and the 75% quantile to assess the risk of a design *ϕ*. The 75%-quantile is the median of the performance distribution truncated below at the median performance. It thus indicates how much worse the loss is expected to get if performing worse than the median.

We found that, in our case studies, the best designs according to the above metrics formed a connected region which we refer to as a cluster. As it is bounded and connected it is natural to find its centroid, which we pick as the single most promising design.

## Code availability

The code is available on GitHub (https://github.com/MichalKobiela/uncertainty-circ-opt). The data and plotting scripts are available on Zenodo (https://zenodo.org/records/13355081).

## Supplementary Material

The Supplementary Materials contain a detailed description of mechanistic models, their parameters, libraries used, and other relevant information.

## Author contributions

MK, DAO, and MUG designed the research. MK implemented the method. All authors analyzed the results. MK wrote the initial manuscript. DAO and MUG edited the manuscript and supervised the research.

## Acknowledgments

This work was supported by the United Kingdom Research and Innovation (grant EP/S02431X/1), UKRI Centre for Doctoral Training in Biomedical AI at the University of Edinburgh, School of Informatics. For the purpose of open access, the author has applied a creative commons attribution (CC BY) licence to any author accepted manuscript version arising.

