## Supplemental Material for "Risk-averse optimization of genetic circuits under uncertainty"

### S1 Models

#### S1 Robust perfect adaptation

##### S1.1 Equations

$$\frac{d[A]}{dt} = \alpha_1 \left( \frac{1}{1 + \left( \frac{K_{TF_{tot}}}{1+\left(\frac{[\text{Input}]}{K_{IR}}\right)} \right)^{n_{RA}}} + \beta_{RA} \right) \left( \frac{[B]^{n_{BA}}}{K_{BA}^{n_{BA}} + [B]^{n_{BA}}} + \beta_{BA} \right) - \gamma_A[A] \tag{S1}$$

$$\frac{d[B]}{dt} = \alpha_2 \left( \frac{1}{1 + \left( \frac{[A]}{K_{AB}} \right)^{n_{AB}}} + \beta_{AB} \right) \left( \frac{[B]^{n_{BB}}}{K_{BB}^{n_{BB}} + [B]^{n_{BB}}} + \beta_{BB} \right) - \gamma_B[B], \tag{S2}$$

##### S1.2 Parameters

For fixed parameters (i.e. neither design nor uncertain parameters) we used the default values that can be found in the table below:

| Parameter | Symbol | Value | Unit |
|---|---|---|---|
| Degradation and dilution rate constant of node A | $\gamma_A$ | 1.0 | $hr^{-1}$ |
| Degradation and dilution rate constant of node B | $\gamma_B$ | 1.0 | $hr^{-1}$ |
| Hill coefficient of Transcription Factor | $n_{ra}$ | 1 | / |
| Hill coefficient of A node | $n_b$ | 1 | / |
| Hill coefficient of B node | $n_{ab}$ | 1 | / |

**Supplementary Table S1** Default parameters for RPA model.

#### S2 Repressilator

##### S2.1 Equations

$$dp_1 = \left(\frac{\alpha}{1+\left(\frac{p_3}{K}\right)^n} - \gamma \cdot p_1\right) dt + \sqrt{\frac{\alpha}{1+\left(\frac{p_3}{K}\right)^n}} dW_{11} + \sqrt{\gamma \cdot p_1} dW_{12} \tag{S3}$$

$$dp_2 = \left(\frac{\alpha}{1+\left(\frac{p_1}{K}\right)^n} - \gamma \cdot p_2\right) dt + \sqrt{\frac{\alpha}{1+\left(\frac{p_1}{K}\right)^n}} dW_{21} + \sqrt{\gamma \cdot p_2} dW_{22} \tag{S4}$$

$$dp_3 = \left(\frac{\alpha}{1+\left(\frac{p_2}{K}\right)^n} - \gamma \cdot p_3\right) dt + \sqrt{\frac{\alpha}{1+\left(\frac{p_2}{K}\right)^n}} dW_{31} + \sqrt{\gamma \cdot p_3} dW_{32} \tag{S5}$$

where $W_{ij}$ represents independent Wiener Processes for $i \in \{1,2,3\}, j \in \{1,2\}$. The Hill coefficient $h$ and degradation rate $\gamma$ are the uncertain parameters $\eta = (h, \gamma)$ while the maximal expression $\alpha$ and Hill constant $K$ are the design parameters $\phi = (\alpha, K)$.

##### S2.2 Parameters

In this case, all the parameters are either the design or uncertain, meaning they are only decided by the algorithm and there are no default values.

#### S2 Libraries used

We used Julia (v1.11.0) language to implement the models using SciML, in particular DifferentialEquation.jl (v7.14.0) [25]. For Bayesian inference, we used Turing.jl (v0.32.3) in case of RPA and KissABC.jl (v3.0.1) for the repressilator. For optimziation (Thompson sampling), we used the automatic differentiation with SciMLSensitivity.jl (v7.68.0) [17] in case of RPA and BayesianOptimziation.jl (v0.2.5) for the Repressilator.

#### S3 Observational noise in RPA

The loss for Thompson sampling including observational noise $\epsilon_i \sim N(0, \sigma^2)$ is:

$$L(\phi) = \mathbb{E}_\Delta \mathbb{E}_{\epsilon_1,\epsilon_2}\left[\frac{(S_1(\phi,\eta)+\epsilon_1-10)^2+(S_2(\phi,\eta,\Delta)+\epsilon_2-10)^2}{2}\right] \tag{S6}$$

$$= \mathbb{E}_\Delta \mathbb{E}_{\epsilon_1,\epsilon_2}\left[\frac{((S_1(\phi,\eta)-10)+\epsilon_1)^2+(S_2(\phi,\eta,\Delta)+\epsilon_2-10)^2}{2}\right] \tag{S7}$$

$$= \mathbb{E}_\Delta \mathbb{E}_{\epsilon_1,\epsilon_2}\left[\frac{(S_1(\phi,\eta)-10)^2+2(S_1(\phi,\eta)-10)\epsilon_1+\epsilon_1^2+(S_2(\phi,\eta,\Delta)+\epsilon_2-10)^2}{2}\right] \tag{S8}$$

$$= \mathbb{E}_\Delta\left[\frac{(S_1(\phi,\eta)-10)^2+(S_1(\phi,\eta)-10)\cdot 0+\sigma^2}{2}\right] + \mathbb{E}_\Delta \mathbb{E}_{\epsilon_2}\left[\frac{(S_2(\phi,\eta,\Delta)+\epsilon_2-10)^2}{2}\right] \tag{S9}$$

$$= \mathbb{E}_\Delta\left[\frac{(S_1(\phi,\eta)-10)^2+\sigma^2}{2}\right] + \mathbb{E}_\Delta \mathbb{E}_{\epsilon_2}\left[\frac{(S_2(\phi,\eta,\Delta)+\epsilon_2-10)^2}{2}\right] \tag{S10}$$

$$= \mathbb{E}_\Delta\left[\frac{(S_1(\phi,\eta)-10)^2+\sigma^2}{2}\right] + \mathbb{E}_\Delta\left[\frac{(S_2(\phi,\eta,\Delta)-10)^2+\sigma^2}{2}\right] \tag{S11}$$

$$= \mathbb{E}_\Delta\left[\frac{(S_1(\phi,\eta)-10)^2+(S_2(\phi,\eta,\Delta)-10)^2}{2}\right] + \sigma^2 \tag{S12}$$

where (5)-(6) is the same process as in (1)-(5) applied to a second term. This shows observational noise independent from $\phi$ simply adds a constant to the original loss. This does not affect the argmin and therefore observational noise can be dropped.

### S4 Strategies to avoid local minima in stochastic gradient optimization of RPA circuit

We used stochastic gradient descent optimization with 10 restarts for each posterior sample. We only kept the best-performing result to avoid local minima. In order to further help the algorithm avoid the local minima, for each subsequent posterior sample, we used the best-performing Thompson sample from the previous run as one of the initial points during multiple restarts.

#### S5 Supplementary Figures

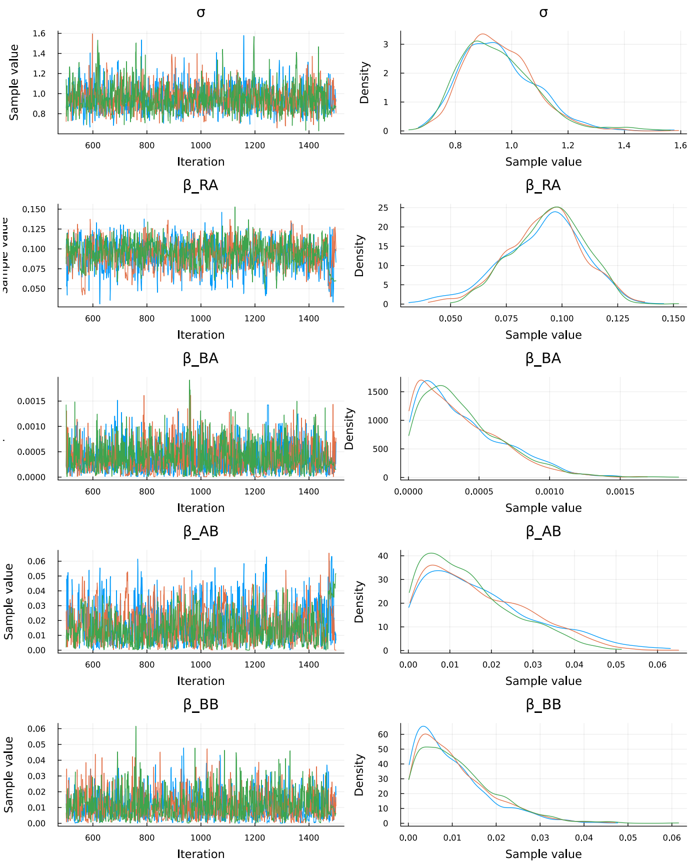


**Supplementary Figure S1** The MCMC chains for the uncertain parameters inference. Three chains were run each resulting in a similar distribution indicating that the method is robust to the initialization point.

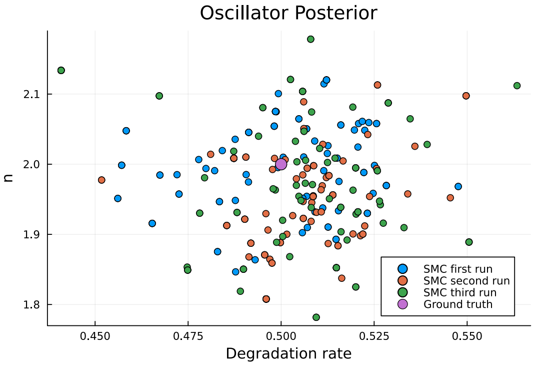


**Supplementary Figure S2** The posterior for the reprissilator over three runs. Note very similar results found each time.

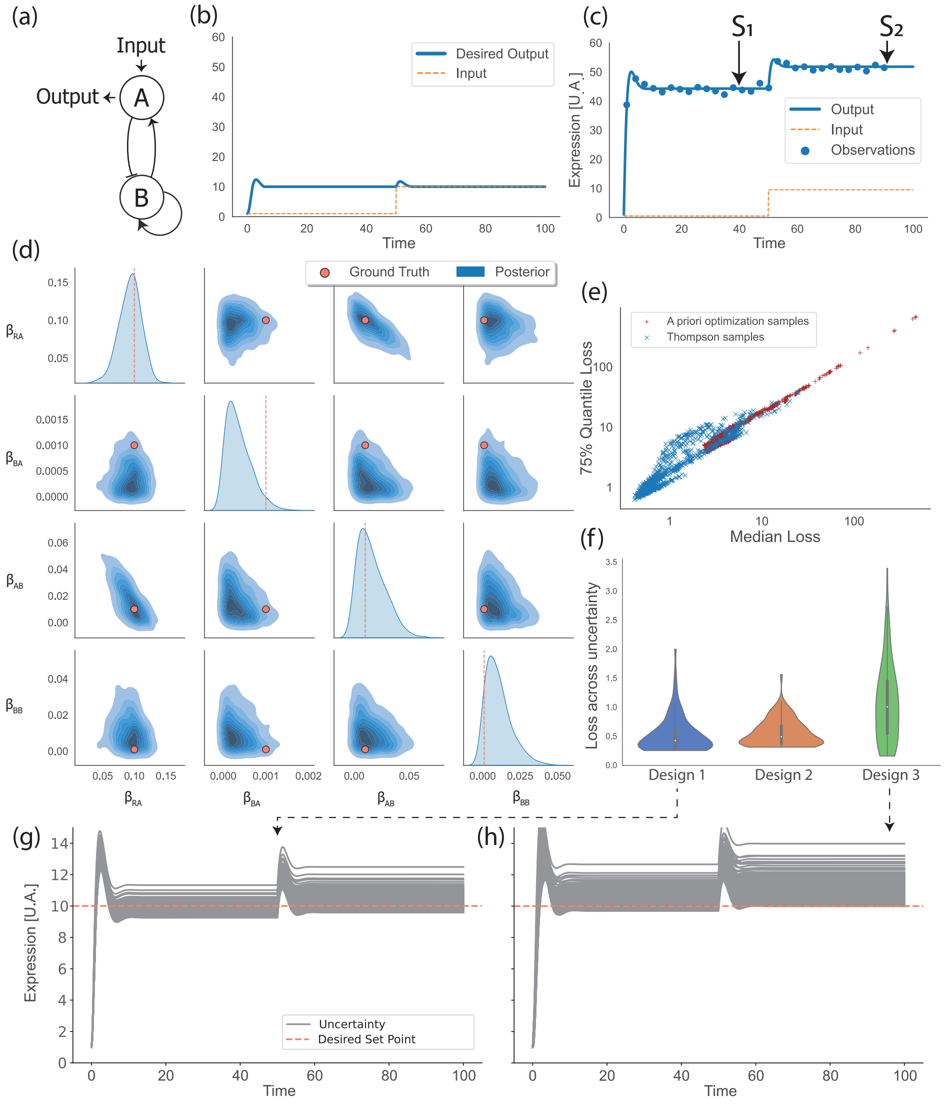


**Supplementary Figure S3** Design of a robust perfect adaptation (RPA) circuit with loss $L(\phi) = \mathbb{E}_\Delta\left[\sqrt{\frac{(S_1(\phi,\eta)-10)^2+(S_2(\phi,\eta,\Delta)-10)^2}{2}}\right]$. (a) Schematic representation of the circuit. (b) The desired behaviour of the circuit adapting to change of input with a set point of 10 U.A. (c) Observed data from a failed design that does not adapt to the change of the input. (d) Posterior over uncertain parameters. (e) Performance of Thompson samples contrasted with *a priori* optimization samples. (f) Performance of three representative designs (Thompson samples). (g) Predictive performance of design 1 (h) Predictive performance of design 3.

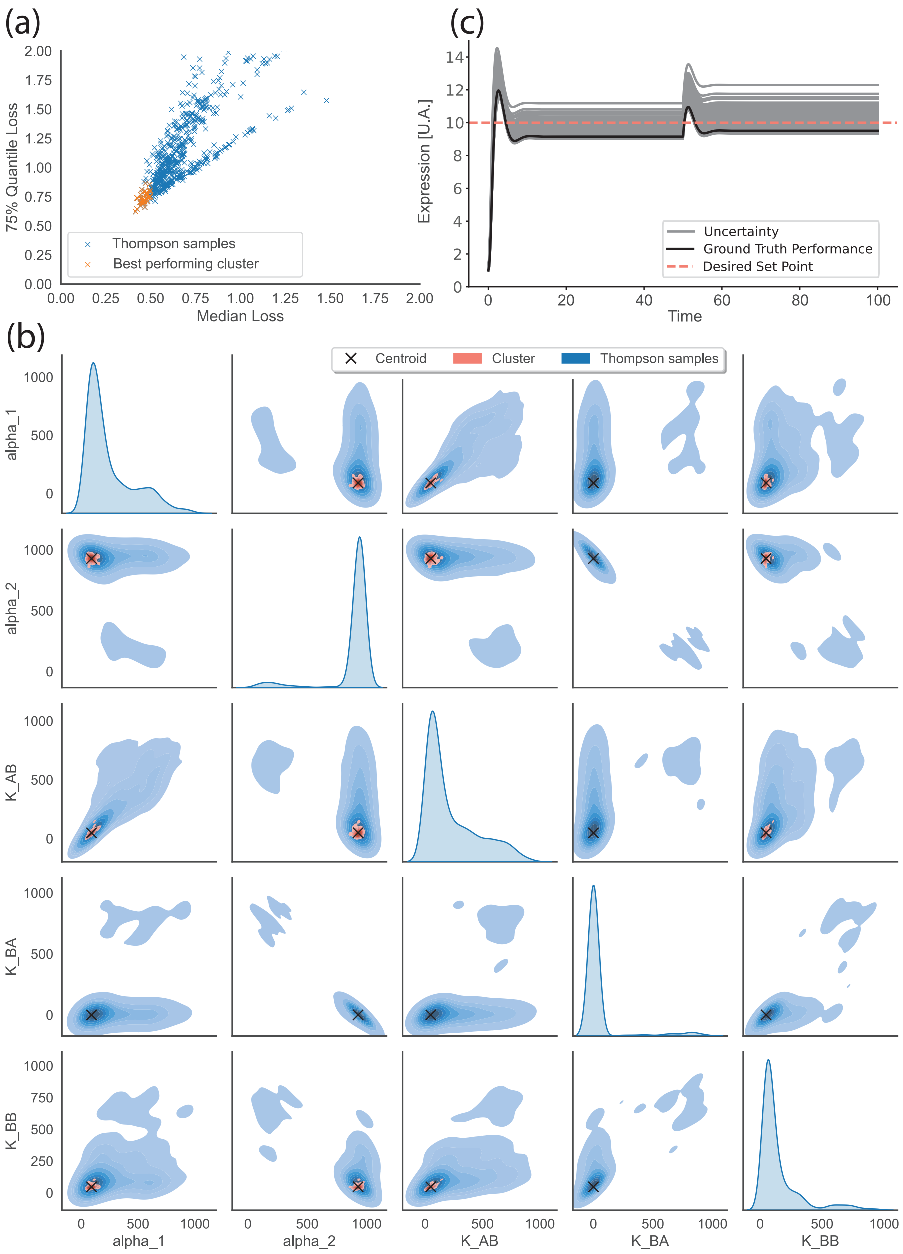


**Supplementary Figure S4** Best performing designs for loss $L(\phi) = \mathbb{E}_\Delta \left[\sqrt{\frac{(S_1(\phi,\eta)-10)^2+(S_2(\phi,\eta,\Delta)-10)^2}{2}}\right]$. Note similar results in terms of suggested designs compared to the loss with no square root. This suggests that using a different loss expressing the same goal still results in successful optimization. (a) Median and 75%-quantile of the loss distribution of each Thompson sample with highlighted best-performing designs. The cut-offs are 0.5 and 0.75 for median and 75%-quantile loss respectively. (b) All the Thompson samples together with the best-performing ones in the design space. The best performing designs turn out to form a cluster. (c) The predictive behaviour of the centroid of the best-performing cluster. A solid black curve shows the performance under ground truth parameters $\eta_{GT}$, while grey trajectories present the performance for multiple posterior samples, thus displaying the epistemic uncertainty.
